# Transcriptome analysis reveals evolutionary co-option of neural development and signaling genes for the wing pigmentation pattern of the polka-dotted fruit fly

**DOI:** 10.1101/2020.01.09.899864

**Authors:** Yuichi Fukutomi, Shu Kondo, Atsushi Toyoda, Shuji Shigenobu, Shigeyuki Koshikawa

## Abstract

How evolutionary novelties have arisen is one of the central questions in evolutionary biology. Pre-existing gene regulatory networks or signaling pathways have been shown to be co-opted for building novel traits in several organisms. However, the structure of entire gene regulatory networks and evolutionary events of gene co-option for emergence of a novel trait are poorly understood. In this study, we used a novel wing pigmentation pattern of the polka-dotted fruit fly, and identified the complete set of genes for pigmentation pattern formation by *de novo* genome sequencing and transcriptome analyses. In pigmentation areas of wings, 151 genes were positively or negatively regulated by *wingless*, a master regulator of wing pigmentation. Genes for neural development, Wnt signaling, Dpp signaling, Zinc finger transcription factors, and effectors (such as enzymes) for melanin pigmentation were included among these 151 genes. None of the known regulatory genes that regulate pigmentation pattern formation in other fruit fly species were included. Our results suggest that the novel pigmentation pattern of the polka-dotted fruit fly emerged through multi-step co-options of multiple gene regulatory networks, signaling pathways, and effector genes, rather than recruitment of one large gene circuit.

## Introduction

How do evolutionary novelties emerge? Researchers have tried to unravel the developmental genetic program underlying traits in order to clarify the origins of evolutionary novelty [1]. Gene regulatory networks for producing novel traits have been supposed to be composed of a combination of genes forming other traits. One of the most significant current discussions regarding the production of evolutionary novelty is how pre-existing regulatory networks were utilized for this production [1,2].

So far, gene regulatory networks or signaling pathways involved in development of novel traits have been scrutinized in several animals. For example, in a horned dung beetle, limb and wing patterning genes are co-opted for horn formation [3,4]. In Nymphalid butterflies, components of the appendage-patterning gene regulatory network, such as *Distal-less*, *wingless*, and *decapentaplegic* signaling, contributed to development of eyespot, another representative novel trait [5–10]. In the fruit fly *Drosophila melanogaster*, it was shown experimentally that the gene regulatory network for larval posterior spiracle development was re-used for the posterior lobe, a novel trait observed in male genitalia [11]. Many studies have shown or suggested which gene regulatory networks or signaling pathways are necessary for, or involved in, development of novel traits. However, the structure of entire gene regulatory networks and evolutionary events of gene co-option for emergence of a novel trait are poorly understood.

Fruit fly species have been used to study regulatory evolution of pigmentation pattern, and provided many examples of mechanisms underlying phenotypic evolution [12–14]. In the polka-dotted fruit fly (*Drosophila guttifera*), which has a novel polka-dotted pigmentation pattern on the wings, a melanin synthesis gene, *yellow*, was expressed in the polka-dotted pattern [15–17]. A Wnt signaling gene, *wingless*, was expressed in the centers of pigmentation areas, and positively regulated the expression of *yellow* through an enhancer (Fig. 1a, c, e) [15,16]. Ectopic expression of *wingless* induced ectopic wing pigmentation (Fig. 1b, d) [16]. The unique expression pattern of *wingless* seemed to be caused by evolutionary gain of novel enhancer activities [18,19]. In *Drosophila melanogaster*, however, there is no pigmentation around crossveins where *wingless* is expressed. If we assume the ancestral species had *wingless* expression and no pigmentation as in *D. melanogaster*, gain of novel expression pattern of *wingless* alone is not sufficient, for emergence of pigmentation pattern [16]. Also, expression of the melanin synthesis gene *yellow* is not sufficient to induce pigmentation in the *Drosophila melanogaster* wing [15,20], indicating that expression changes of multiple genes were required for the evolution of pigmentation. Therefore, exploring the complete set of genes involved in pigmentation pattern formation, which include both regulatory and effector genes, is necessary for understanding emergence of the novel wing pigmentation pattern.

In this study, we identified the complete set of genes for pigmentation formation, by *de novo* genome sequencing and two successive transcriptome analyses by Quartz-Seq, a highly sensitive method of RNA sequencing. In the first transcriptome analysis, we compared gene expression patterns between pigmentation areas and an unpigmented area and searched for differentially expressed genes (DEGs). In the second transcriptome analysis, we tested whether those DEGs were regulated by *wingless*, the master control gene for wing pigmentation.

**Fig. 1.**
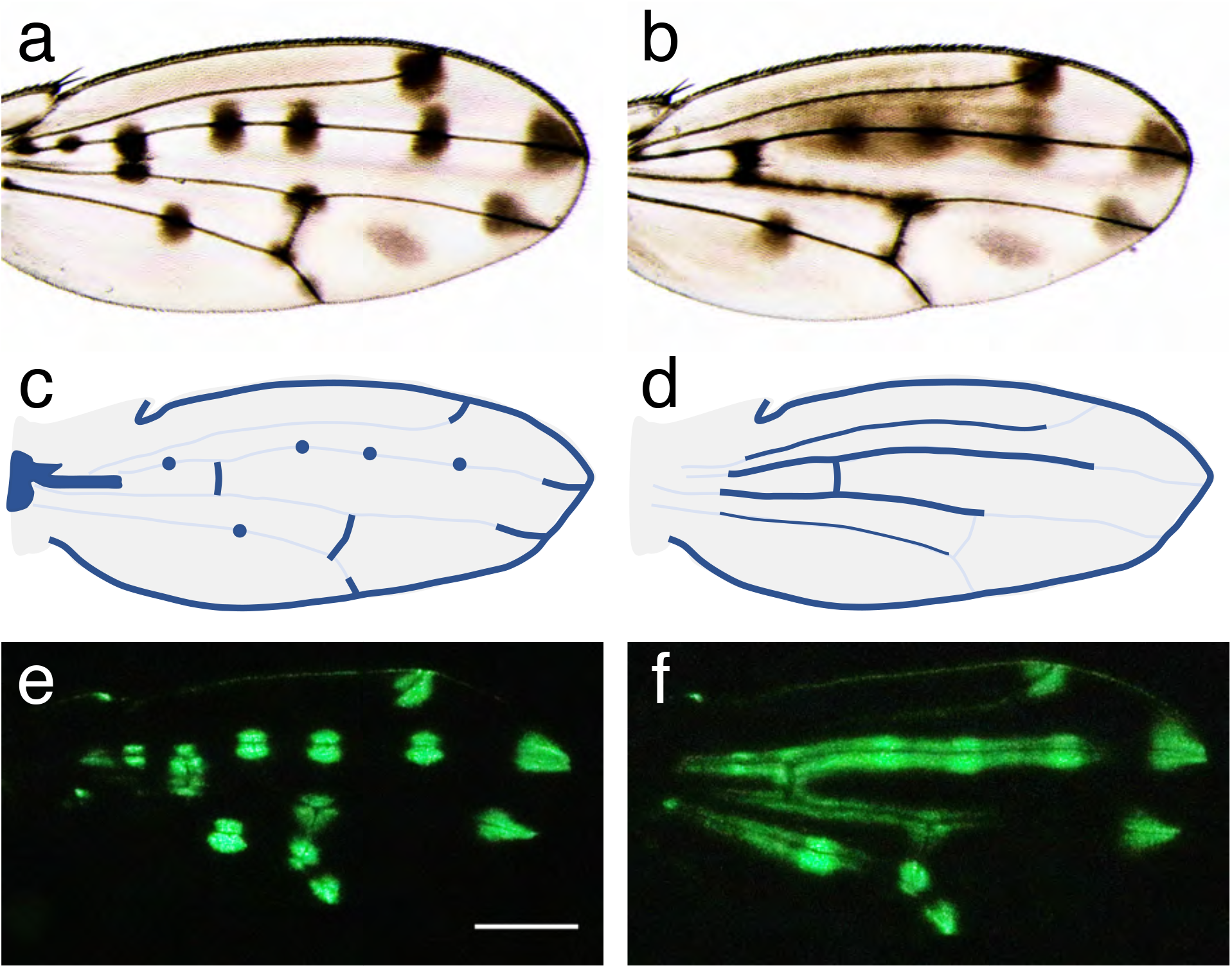
Wings of *Drosophila guttifera* and the expression pattern of *wingless*. **a**: A wing of *Drosophila guttifera*. This picture shows wild type pigmentation pattern. **b**: The wing pigmentation pattern of an individual in which *wingless* is ectopically expressed. **c**: The expression pattern of *wingless* (blue) in a wing of a wild type individual. **d**: The position where *wingless* is ectopically expressed (blue) in the fly shown in b (Modified from Werner et al. 2010). **e**: The expression pattern of EGFP driven by an enhancer of *yellow*. In this individual, the expression pattern of *wingless* is the same as wild type. **f**: The expression pattern of EGFP driven by an enhancer of *yellow*. in an individual with ectopic *wingless* expression shown in d. The scale bar indicates 250 μm

## Results

### Genes expressed in the polka-dotted pattern

We compared gene expression patterns between pigmentation areas and an unpigmented area, and searched differentially expressed genes (DEGs). These areas can be distinguished by GFP label using a transgenic line which carries *eGFP* connected with an enhancer of *yellow* (Werner et al. 2010) (Fig. 1e, f). We identified genes upregulated or downregulated commonly in Area 1 (pigmentation area around a campaniform sensillum, Fig. S1a) and Area 2 (vein tip, Fig. S1a), compared with Area 3 (unpigmented, Fig. S1b). Comparison of the gene expression between Area 1 and Area 3 showed that 2333 genes were differentially expressed. Among them, 1390 genes were upregulated (Fig. 2a) and 943 genes were downregulated (Fig. 2b) in Area 1 in comparison to Area 3. 2582 genes were differentially expressed between Area 2 and Area 3. Among them, 1593 genes were upregulated (Fig. 2a) and 989 genes were downregulated (Fig. 2b) in Area 2 in comparison to Area 3. Integrating these data, the number of common DEGs was 1035. Among them, 615 genes were upregulated both in Area 1 and Area 2 (Fig. 2a), while 420 genes were downregulated in Area 1 and Area 2 (Fig. 2b). Consistent with previously reported findings, *wingless* and *yellow* were expressed in the pigmentation areas [16,18], indicating the high sensitivity and accuracy of the present method of analysis. *wingless* and *yellow* were included in the 615 commonly upregulated DEGs (Fig. 3, Table S1, Table S2).

**Fig. 2.**
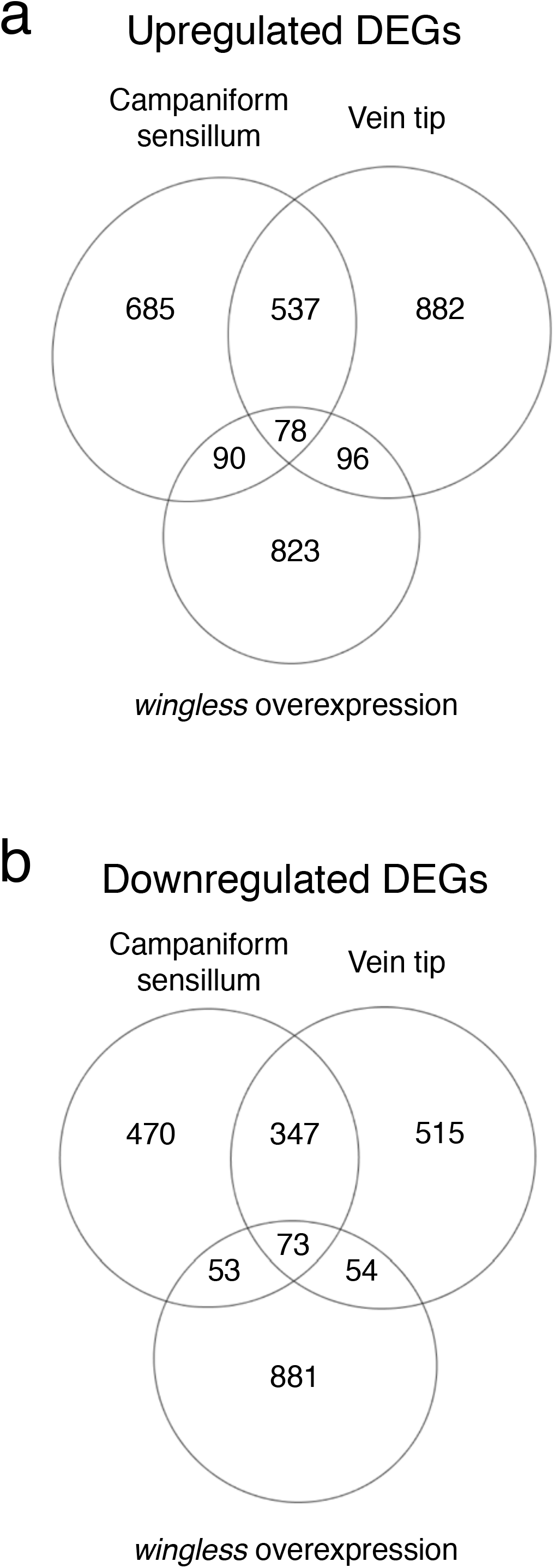
The number of differentially expressed genes (DEGs) detected in transcriptome analyses. **a**: The number of DEGs upregulated at pigmentation areas and an area where *wingless* is ectopically expressed. **b**: The number of DEGs upregulated at the pigmentation areas and an area where *wingless* is ectopically expressed. The circle labeled with “Campaniform sensillum” indicates the result from the comparison of transcriptome between Area 1 (pigmentation area around campaniform sensillum) and Area 3 (unpigmented). The circle labeled with “Vein tip” indicates the result from the comparison of transcriptome between Area 2 (a pigmentation area at the tip of 3rd longitudinal vein) and Area 3. The circle labeled with “*wingless* overexpression” indicates the number of genes differentially expressed where *wingless* is ectopically expressed.

**Fig. 3.**
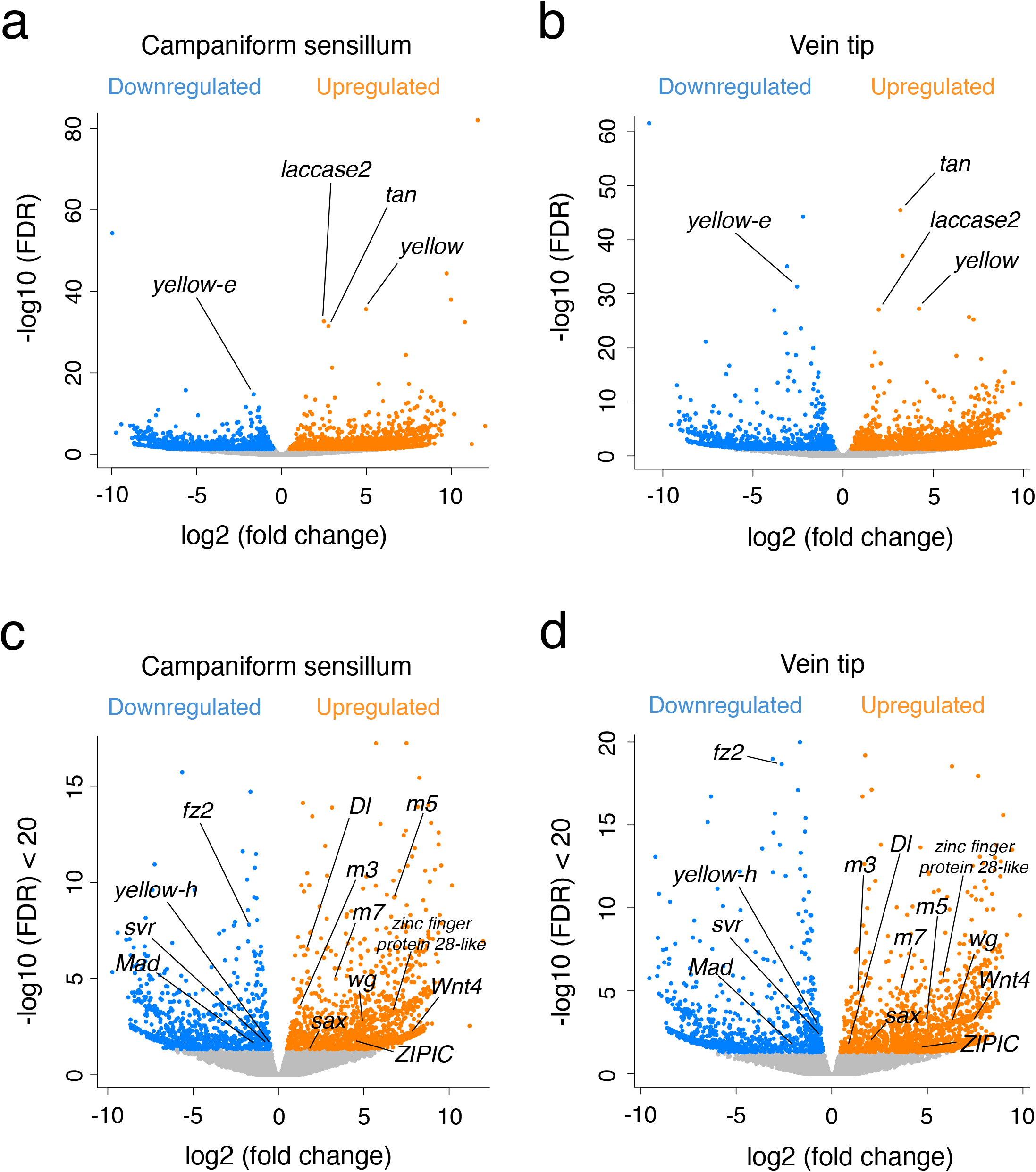
Volcano plots of the results from the first transcriptome analysis. The horizontal axis indicates fold changes and the vertical axis indicates significance calculated as FDR with edgeR. Orange points indicate upregulated DEGs and blue points indicate downregulated DEGs. **a**: This plot shows genes differentially expressed in Area 1 (a pigmentation area around a campaniform sensillum) compared with gene expression of Area 3 (unpigmented). **b**: This plot shows genes differentially expressed in Area 2 (a pigmentation area at the tip of 3rd longitudinal vein) compared with gene expression of Area 3. **c**: Genes with −log 10 (FDR) < 20 are extracted from a. **d**: Genes with −log 10 (FDR) < 20 are extracted from b. In c and d, *m3*, *m5*, and *m7* indicate *E(spl)m3-HLH*, *E(spl)m5-HLH*, and *E(spl)m7-HLH* respectively.

### Pigmentation pattern-associated genes regulated by *wingless*

Because ectopic expression of *wingless* is known to induce pigmentation, genes sufficient for pigmentation formation in wings must be included in the gene network downstream of *wingless*. To identify the genes that are under the control of *wingless*, we identified genes upregulated or downregulated when *wingless* was ectopically expressed. Among the 615 common upregulated (Area 1 and 2) DEGs, 78 genes were upregulated by ectopic expression of *wingless* (Fig. 2a). In 420 common downregulated (Area 1 and 2) DEGs, *wingless* downregulated 73 genes (Fig. 2b). In total, 151 genes associated with the pigmentation pattern were regulated by *wingless* gene. These 151 genes were blasted against the protein database of *Drosophila melanogaster* and 131 genes were annotated. For these 131 genes, enrichment analysis with DAVID resulted 14 functional annotation clusters, and 6 of which were significant (Table S1). In the most significant cluster, Gene Ontology (GO) terms “Glycoprotein”, “Plasma membrane”, “Disulfide bond”, “Signal peptide” and “Receptor” were included (Table 3). GO terms such as “cuticle pigmentation” and “melanin biosynthetic process” were included in the 3rd significant cluster. 20 genes that could not be annotated were reanalyzed with Blast2GO. Four genes were annotated and remaining 16 genes did not match to any gene in the database.

Among pigmentation pattern associated genes regulated by *wingless*, six genes can be categorized as melanin synthesis-related genes [21–23]. Among them, *yellow*, *laccase2*, and *tan* were upregulated in the pigmentation areas (Fig. 3a, b, Table S1, Table S2) and also upregulated by *wingless* (Fig. 4a, Table S4). *yellow-e*, *yellow-h*, and *silver* (*svr*) were downregulated both in the pigmentation areas (Fig. 3a, b, c, d, Table S1, Table S2) and in the area where *wingless* was ectopically expressed (Fig. 4a, b, Table S3).

**Fig. 4.**
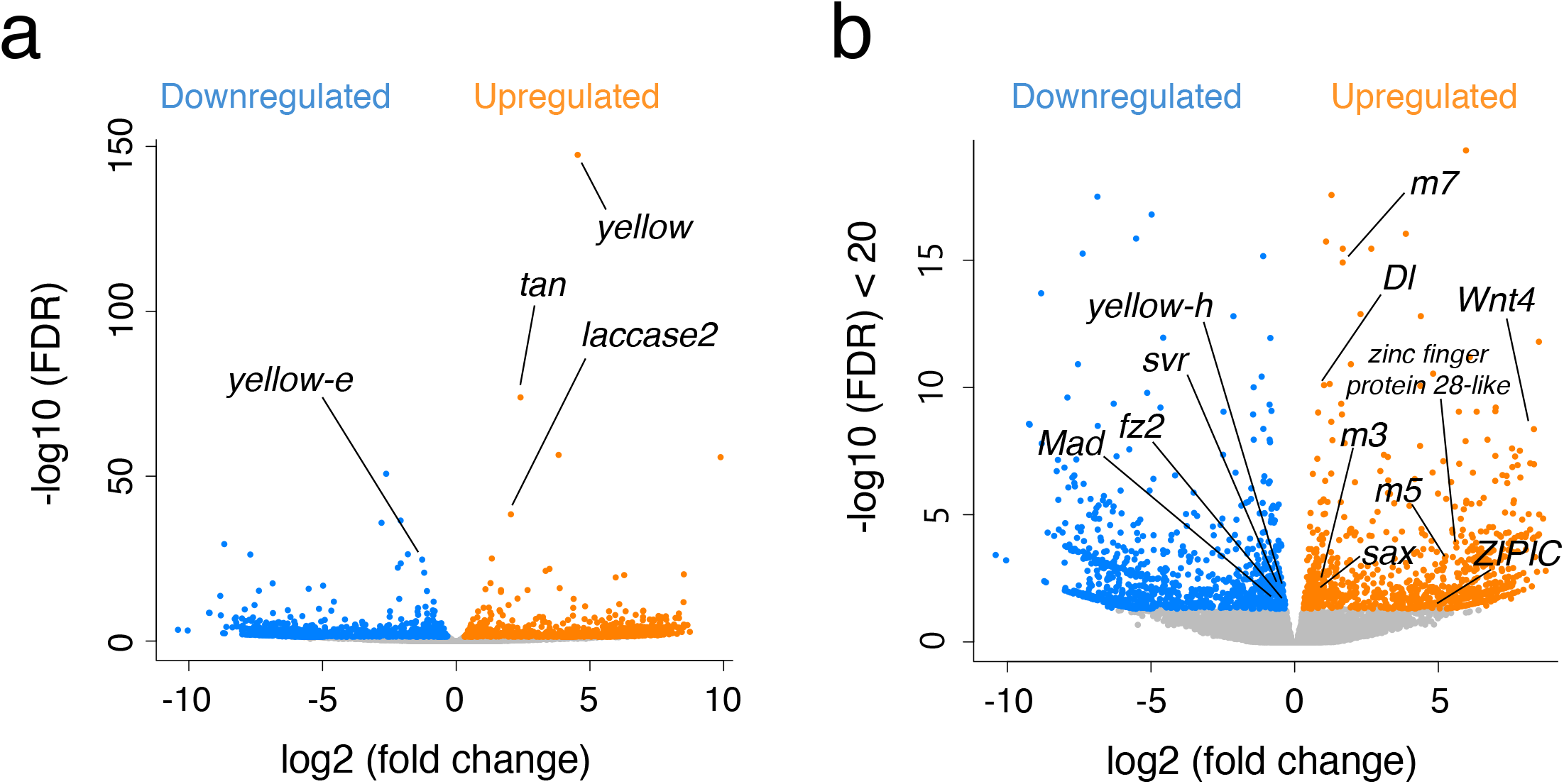
Volcano plots of the results from the second transcriptome analysis. The horizontal axis indicates fold changes and the vertical axis indicates significance calculated as FDR with edgeR. Orange points indicate DEGs upregulated and blue points indicate downregulated DEGs. **a**: This plot shows genes differentially expressed where *wingless* is ectopically expressed. **b**: Genes with −log 10 (FDR) < 20 are extracted from **a**.

Regulatory genes, such as transcription factors and genes involved in signaling pathways are important to understand the regulatory network controlling the pigmentation pattern. Six transcription factors were associated with pigmentation and regulated by *wingless*. *Zinc-finger protein interacting with CP190* (*ZIPIC*), *zinc finger protein 28-like*, and Enhancer of split complex genes such as *E(spl)m3-HLH*, *E(spl)m5-HLH*, and *E(spl)m7-HLH* were upregulated in the pigmentation areas (Fig. 3c, d, Table S1, Table S2) and by *wingless* (Fig. 4b, Table S3). *Mothers against dpp* (*Mad*) was downregulated in the pigmentation areas and by *wingless*. Two signal ligands, *Delta* (*Dl*) and *Wnt oncogene analog 4* (*Wnt4*) were upregulated in the pigmentation areas and by *wingless* (Fig. 3c, d, Fig. 4b, Table S1, Table S2, Table S3). As receptors of ligands, *saxophone* (*sax*) was upregulated and *frizzled 2* (*fz2*) was downregulated in the pigmentation areas and by *wingless* (Fig. 3c, d, Fig. 4b, Table S1, Table S2, Table S3). *wingless* itself was not detected in DEG analysis with ectopic *wingless* expression, which is reasonable in our experimental design. Our transgenic line ectopically drove the *wingless* gene originated from *D. melanogaster* [16] and its transcripts were not mapped on *D. guttifera* genome in the analysis.

## Discussion

### A large number of genes were specifically regulated in the area of pigmentation formation

Transcriptome analyses revealed that a surprisingly large number of genes were specifically regulated in the area of pigmentation formation: 78 genes were upregulated commonly in the pigmentation area and by *wingless*, and 73 genes were downregulated commonly in the pigmentation area and by *wingless*. In the butterfly *Bicyclus anynana*, 132 genes were upregulated and 54 genes were downregulated in relation to eye spot formation [24]. In comparison with these numbers of genes in butterflies, the number of genes identified by our results seem reasonable. However, *Drosophila* pigmentation has been thought to be a simple trait, and researchers have tried to explain the evolution of pigmentation by changes of expression of a small number of genes [15,25–27]. In the abdominal tergite of *Drosophila melanogaster*, the combination of *ebony* mutation and ectopic expression of the *yellow* gene can induce ectopic pigmentation [28]. In *D. melanogaster* wings, however, the same combination resulted in scarcely any ectopic pigmentation [15,20]. Those findings are consistent with those of the present study, in which we found many genes that were specifically regulated in the areas of pigmentation formation. This also suggests that experimental reproduction of the gain of pigmentation patterns through overexpression of genes is not trivial.

Although ectopic expression of *wingless* can induce pigmentation, the intensity of induced pigmentation is weaker than that of the natural spotted pigmentation. Five hundred thirty-seven genes were upregulated and 347 genes were downregulated commonly in Area 1 and Area 2 but not regulated by *wingless* (Fig. 2). Thus, they were not essential to make pigmentation, but might have supplemental roles, or they might be unrelated to pigmentation but have a structural role unique to the pigmented area.

### Known gene regulatory networks of *Drosophila* pigmentation were not responsible for *D. guttifera* wing pigmentation

Regulation of pigmentation has been studied in multiple *Drosophila* models. The best-studied case was abdominal pigmentation in *D. melanogaster*. Male-specific pigmentation was controlled positively by *Abd-B* and negatively by *bab* genes, and pigmentation common to the two sexes was positively controlled by *omb* [29]. In *D. biarmipes* wings, the pigmentation is controlled positively by *Dll*, and negatively by *en* [13,15]. Neither of these genes was included in DEGs in our analysis. Thus, regulatory mechanisms of pigmentation in *D. guttifera* are not a simple co-option of the known gene regulatory network of pigmentation. Then, what kinds of genes are responsible for the emergence of the novel pigmentation pattern in *D. guttifera*?

### Neural development genes and other signaling genes were co-opted for the pigmentation areas

We identified 10 regulatory genes (transcription factor genes and signaling factor genes) that were regulated by *wingless* in the pigmentation areas. Among them, four (*Dl*, *E(spl)m3-HLH, E(spl)m5-HLH* and *E(spl)m7-HLH*) genes belonged to neurogenesis genes of GO terms. Because Area 2 (vein tip) did not include any tissue of neural origin, these genes were considered to be expressed in epidermal cells, which is the only cell type present in Area 2. In wing discs of *D. melanogaster*, *Dl* and *E(spl)* complex genes are involved in neurogenesis mediated by Notch signaling [30]. During this process, the expression of *wingless* is necessary and sufficient for the expression of *Dl* [31]. The gene regulatory network of neurogenesis, including *Dl* and *E(spl)* complex, might be co-opted to wing pigmentation of *D. guttifera*, probably in relation to its regulatory gene, *wingless* (Fig. 5).

**Fig. 5.**
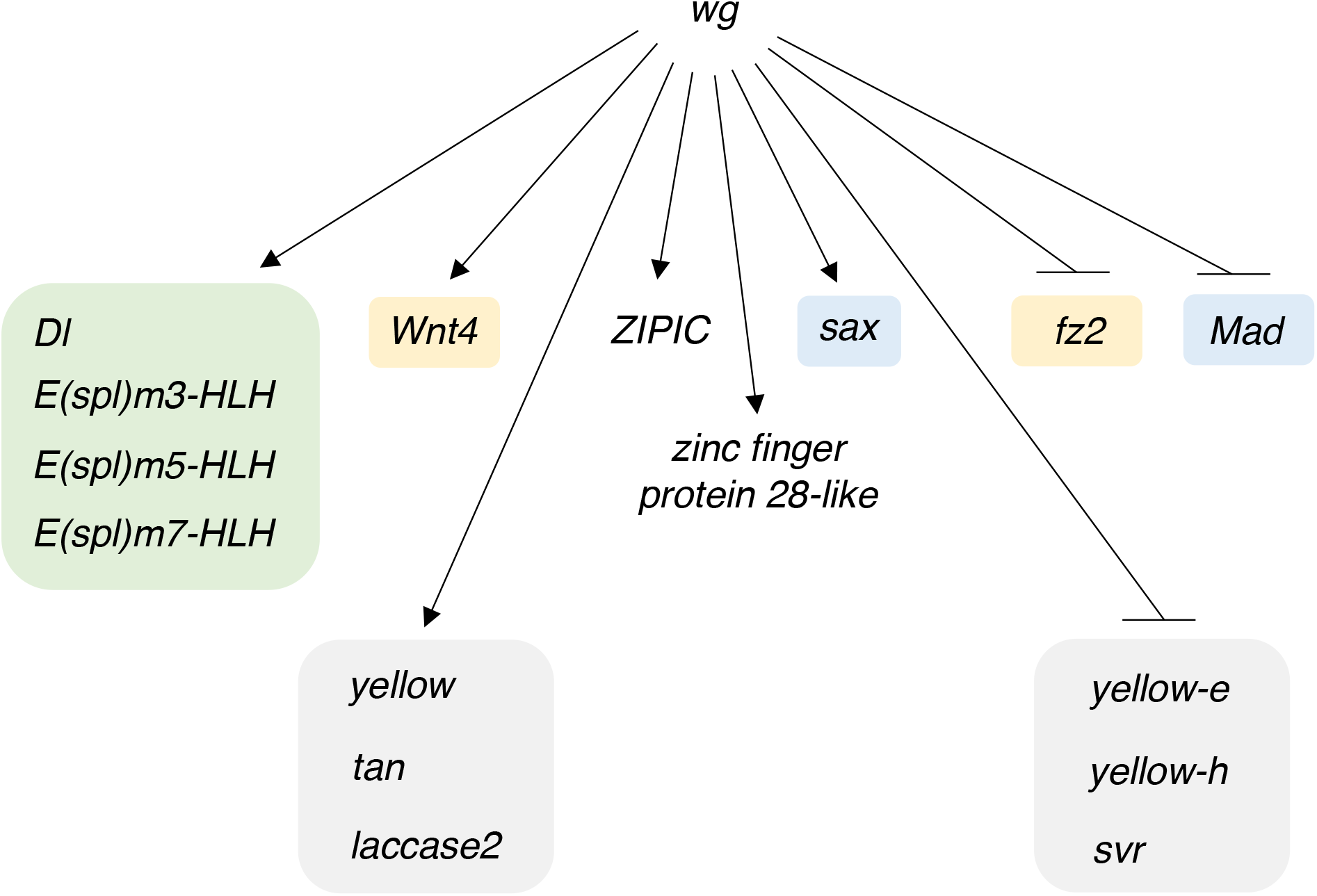
The putative gene regulatory network for formation of the novel wing pigmentation pattern. Green box: Notch signaling gene, grey boxes: melanin synthesis genes, yellow boxes: Wnt signaling genes, and blue boxes: Dpp signaling genes.

Dpp and Hh signaling can work cooperatively or antagonistically with Wingless signaling in various aspects of *Drosophila* development [32,33]. In the present study, genes involved in Dpp signaling, *saxophone* (*sax*) and *Mothers against dpp (Mad)*, were regulated by *wingless* in the pigmentation areas*. hh* and *patched* of the Hh signaling pathway were up-regulated in the area of pigmentation, but not regulated by *wingless* (Table S4). Hh signaling may play a role in the control of pigmentation by affecting Wingless signaling.

Wnt signaling genes *fz2* and *Wnt4* were regulated in the pigmentation areas and by *wingless* (Fig. 5). Downregulation of *fz2* in the pigmentation areas of *D. guttifera* might contribute to achieving the proper gradient of Wingless protein, as it is known to do in *D. melanogaster* wing discs [34]. *Wnt4* was expressed in the area surrounding the *wingless* expression region in wings of *D. guttifera* [18], suggesting that it might play a role in pigmentation pattern formation.

Two genes encoding zinc finger transcription factors, *ZIPIC* [35] and *zinc finger protein 28-like*, were also upregulated in the pigmentation areas and by *wingless*, suggesting that they may play a role in regulation of pigmentation (Fig. 5).

### Known functions of pigmentation genes regulated by *wingless*

In *D. melanogaster*, *yellow*, *tan*, and *laccase2* are known to be effector genes for pigmentation and have been proven to promote melanin pigmentation in *D. melanogaster* [20,28,36]. Association of these genes with melanin pigmentation patterns was also reported in other insects [22,37,38]. Therefore it was not surprising to detect these three genes in the present study. The *svr* gene encodes a carboxypeptidase and is involved in metabolism of N-acetyl dopamine (NADA) in *D. melanogaster* [21,39]. As the expression of *svr* was downregulated in the area of pigmentation of *D. guttifera*, inhibiting the metabolism of NADA might contribute to wing pigmentation. Expression of Yellow family protein genes such as *yellow-e* and *yellow-h* was also downregulated in the pigmentation areas on the wings of *D. guttifera*, but the molecular functions of these genes have not been identified in any insect. In wings of the butterflies (*Vanessa caudui* and *Heliconius* spp.), another Yellow family protein gene, *yellow-d*, was upregulated at red pigmentation areas compared with black pigmentation areas [38,40]. Our results together with these previously reported studies suggest that these Yellow protein family genes might play a role in inhibiting melanin pigmentation.

### Evolutionary scenario of pigmentation pattern evolution

The genes identified in the present study, especially genes regulated by *wingless*, were highly likely to have been co-opted for the pigmentation formation. This raises the question: What kind of genetic change enabled the evolution of pigmentation patterns? There is an ongoing discussion about whether a large gene circuit is recruited or many genes are individually recruited to form a circuit during the evolution of a novel trait [41,42]. Examples of co-option of pre-existing circuits consisting of multiple regulatory genes are known from animals and plants [11,24,43–46]. In the present case, we can ask whether the large gene network for pigmentation regulated by a master control gene, *wingless*, was co-opted all at the same time, or whether individual genes were co-opted one by one to form the current network. To our knowledge, there is no functional evidence of a pigmentation pattern mainly regulated by *wingless* in other *Drosophila* species. Therefore, a “one by one model” seems reasonable to explain our experimental data. The current network could consist of multiple subnetworks. For example, the circuit of neural developmental genes (*Dl*, *E(spl)m3-HLH, E(spl)m5-HLH* and *E(spl)m7-HLH*) could be co-opted all at the same time from neural development to regulation of pigmentation.

Taking our findings altogether, we conclude that the novel pigmentation pattern of *D. guttifera* could have been caused by multi-step co-options of gene circuits, regulatory genes and effector genes. To test this scenario, we will have to compare multiple species with and without pigmentation, as well as to test individual gene functions in *D. guttifera*. These investigations will further our understanding of the evolution of pigmentation pattern formation in *D. guttifera*, as an example of the emergence of evolutionary novelties.

## Materials and methods

### Flies

*Drosophila guttifera* is a North American species that belongs to (or is closely related to) the *quinaria* group of subgenus *Drosophila* [47,48]. The inbred line (A5) was made by ten successive sibling crosses of a wildtype (stock no. 15130-1971.10) obtained from the *Drosophila* Species Stock Center at the University of California, San Diego. Two lines (transgenic lines No. 1 and No.2) of *D. guttifera* were used for transcriptome analyses. Transgenic line No. 1 was established by five successive backcrosses (introgression) of a transgenic line that carries *nuclear eGFP* connected with a *yellow* enhancer (*vein spot* CRE-*nuclear eGFP*, gut 1c+R GFP #12) [16] with the A5 inbred line. Transgenic line No. 2 was established by two successive backcrosses of a UAS-*wg* line [16] with transgenic line No. 1. These backcrosses aimed to unify the genetic backgrounds to improve the mapping efficiency in the transcriptome analyses. Flies were reared with standard cornmeal/sugar/yeast/agar food at 25 °C [49].

#### (b) Genome sequencing and gene prediction

Genomic DNA was extracted from adults of the inbred line (A5) with a Gentra Puregene Tissue Kit (Qiagen). A paired-end library with an average insert size of 450 bp was constructed with a TruSeq DNA PCR-free Library Prep Kit (Illumina) and two different mate-pair libraries (3 kb and 5 kb) were prepared with a Nextera Mate Pair Library Prep Kit (Illumina). Sequencing libraries were run on the Illumina Hiseq 2500 sequencer with a read length of 150 bp. Paired-end and mate pair reads were *de novo* assembled using Platanus v1.2.1.1 [50] after removal of adaptors and error correction with SOAPec [51] (Table 1, Table 2). The assembly sequences had 767 scaffolds with a total length of 168.4 Mb and a scaffold N50 length of 1.8 Mb.

**Table 1.**
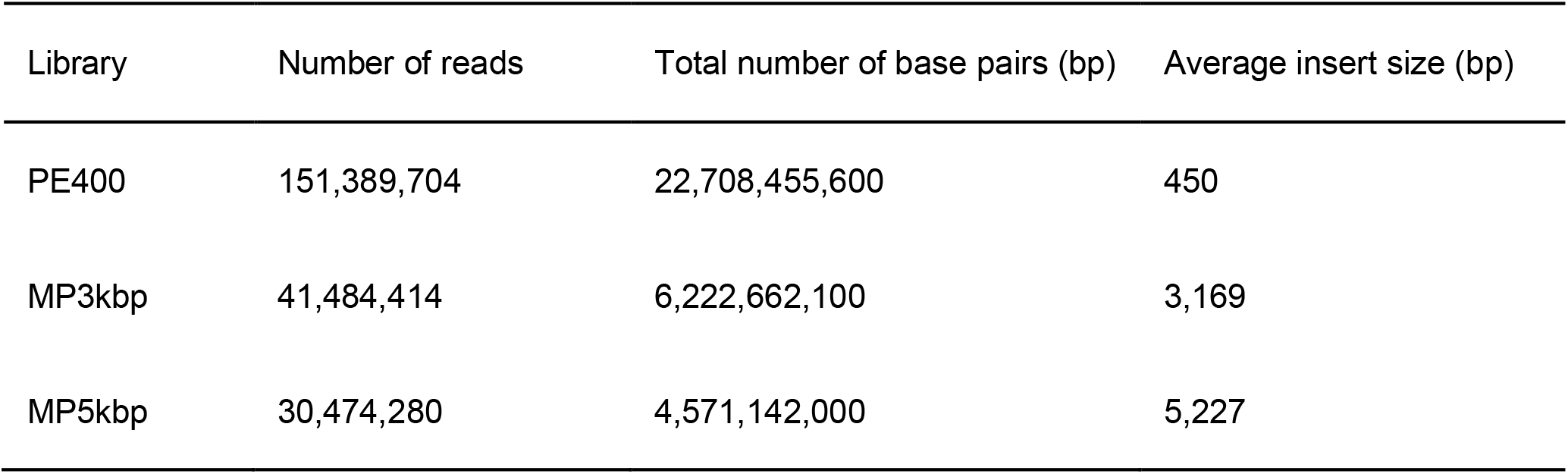
The data of constructed libraries of *D. guttifera*. The data was obtained by genome sequencing with Hiseq 2500. PE400, MP3kbp, and MP5kbp respectively indicate the paired-end library, the mate-pair library (3kb), and the mate-pair library (5kb).

**Table 2.**
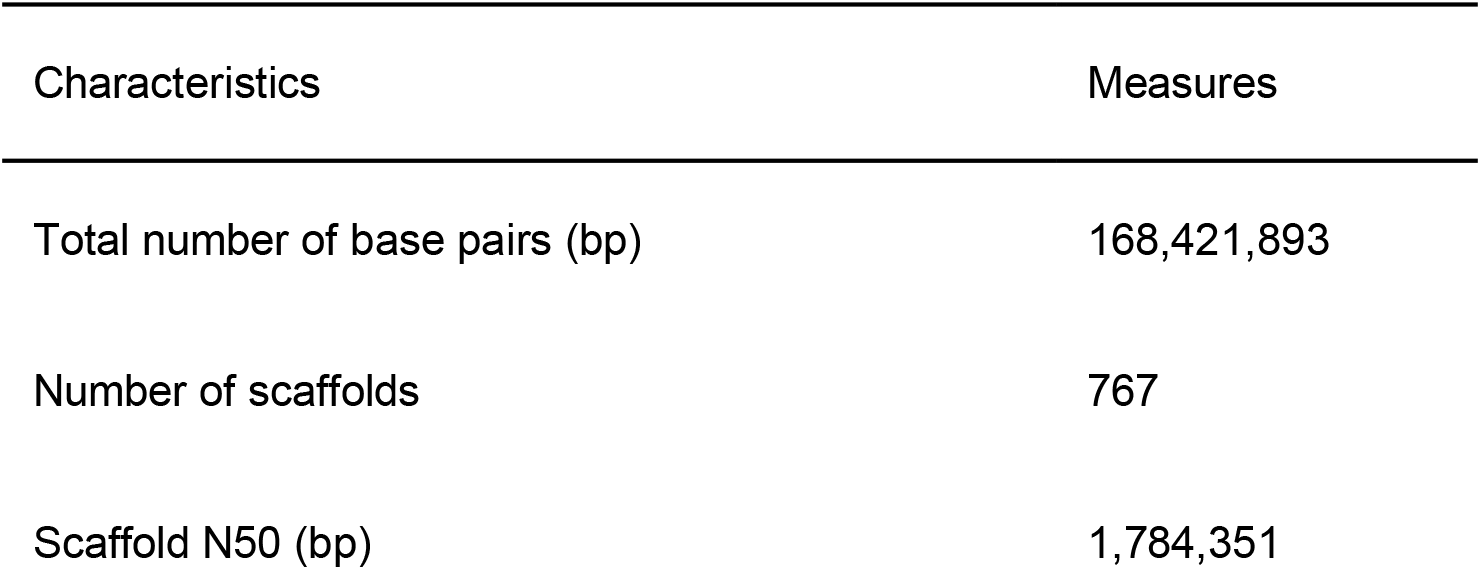
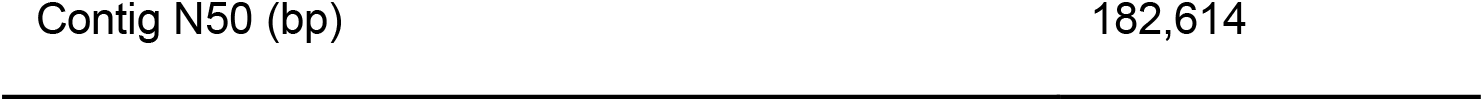
Characteristics of the *D. guttifera* genome sequence obtained by *de novo* assembly.

**Table 3.**
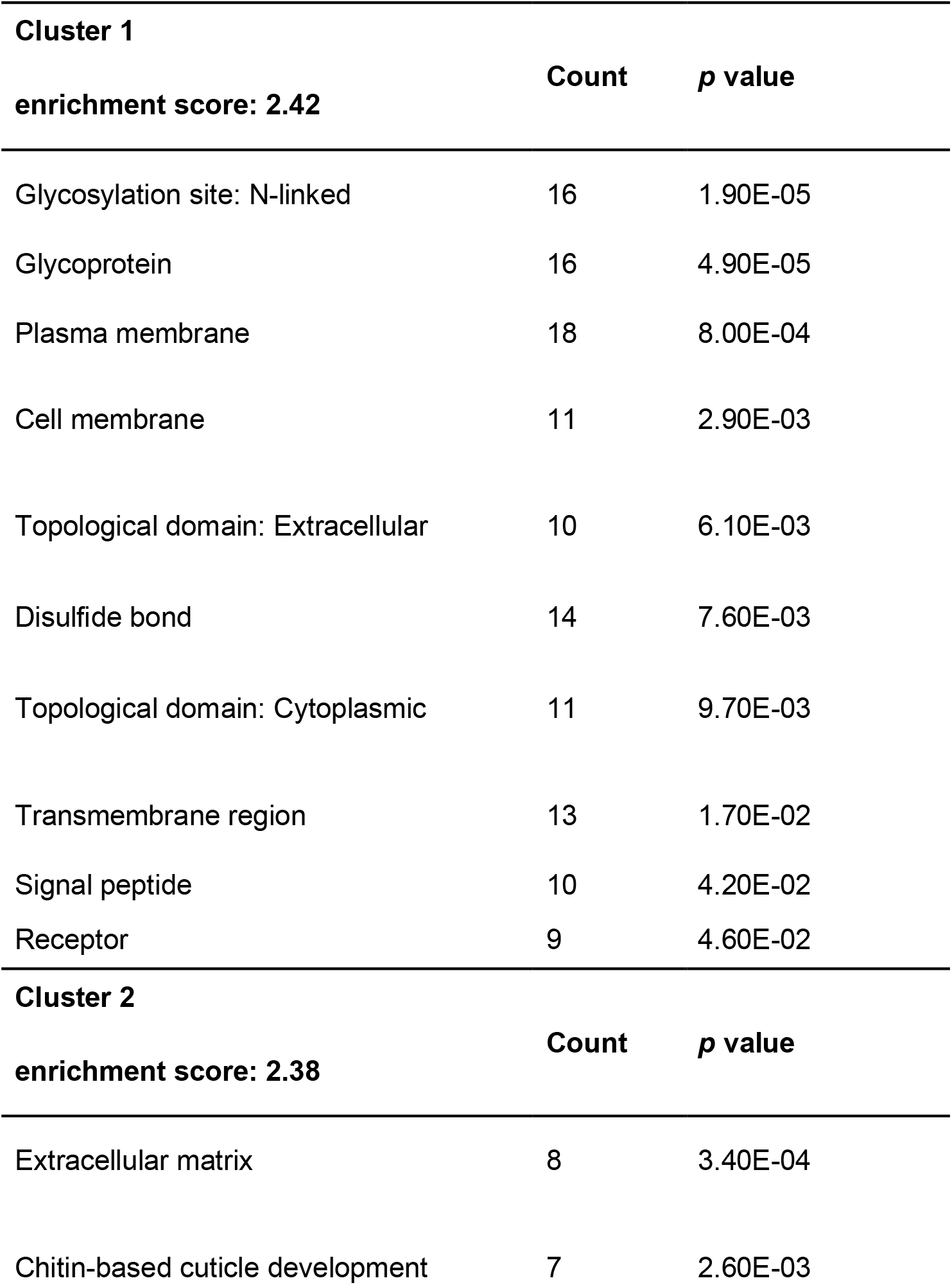
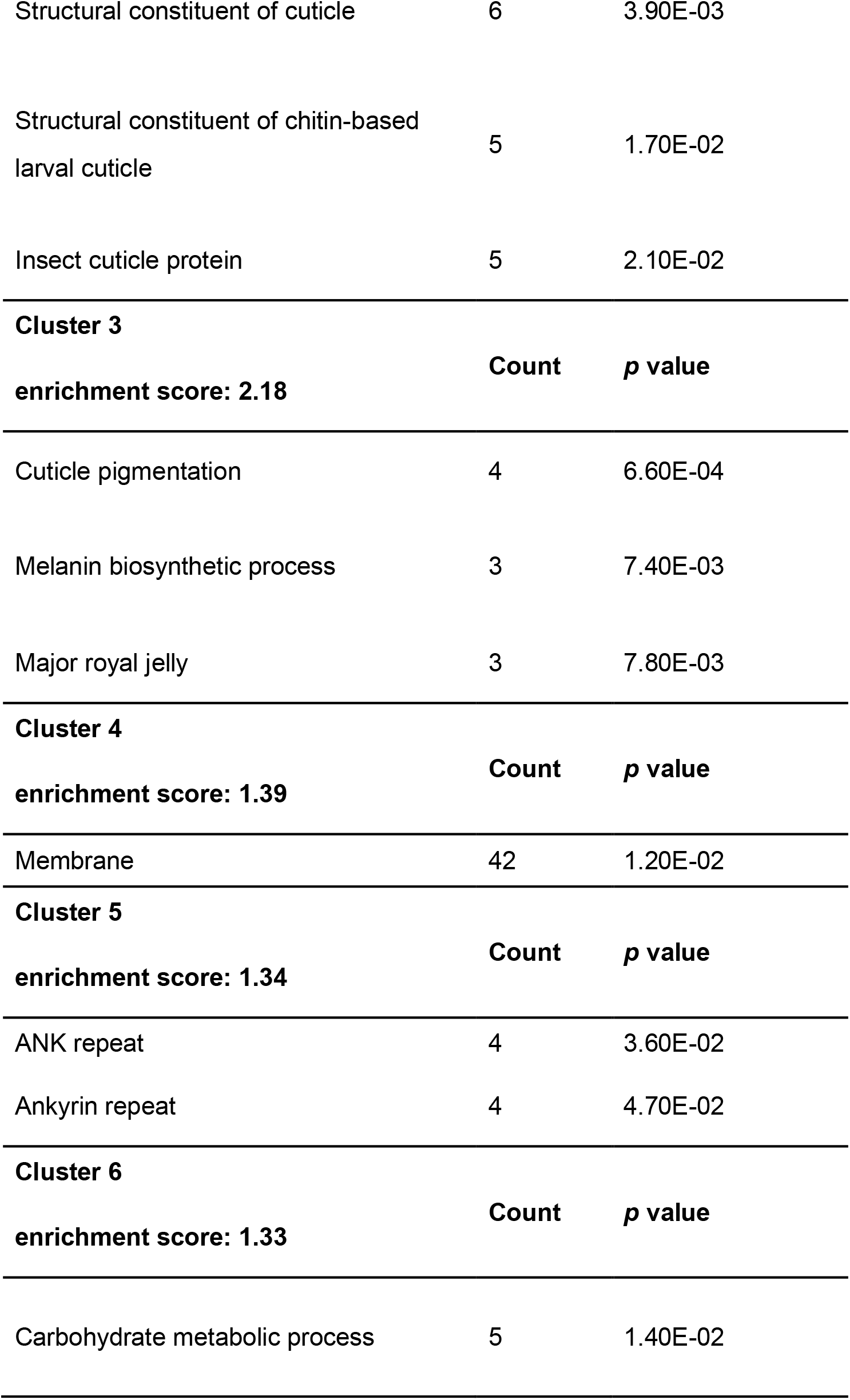
The result from enrichment analysis with DAVID, showing clusters with enrichment score > 1.3 and Gene ontology terms with *p* value < 0.05 in each cluster (Count indicates the number of genes for each Gene ontology term).

| **Cluster 1** |  |  |
| --- | --- | --- |
| **enrichment score: 2.42** | **Count** | **$p$ value** |
| Glycosylation site: N-linked | 16 | 1.90E-05 |
| Glycoprotein | 16 | 4.90E-05 |
| Plasma membrane | 18 | 8.00E-04 |
| Cell membrane | 11 | 2.90E-03 |
| Topological domain: Extracellular | 10 | 6.10E-03 |
| Disulfide bond | 14 | 7.60E-03 |
| Topological domain: Cytoplasmic | 11 | 9.70E-03 |
| Transmembrane region | 13 | 1.70E-02 |
| Signal peptide | 10 | 4.20E-02 |
| Receptor | 9 | 4.60E-02 |
| **Cluster 2** |  |  |
| **enrichment score: 2.38** | **Count** | **$p$ value** |
| Extracellular matrix | 8 | 3.40E-04 |
| Chitin-based cuticle development | 7 | 2.60E-03 |
| Structural constituent of cuticle | 6 | 3.90E-03 |
| Structural constituent of chitin-based larval cuticle | 5 | 1.70E-02 |
| Insect cuticle protein | 5 | 2.10E-02 |

Cluster 3 **enrichment score: 2.18**
|  | Count | *p* value |
| --- | --- | --- |
| Cuticle pigmentation | 4 | 6.60E-04 |
| Melanin biosynthetic process | 3 | 7.40E-03 |
| Major royal jelly | 3 | 7.80E-03 |

Cluster 4 **enrichment score: 1.39**
|  | Count | *p* value |
| --- | --- | --- |
| Membrane | 42 | 1.20E-02 |

Cluster 5 **enrichment score: 1.34**
|  | Count | *p* value |
| --- | --- | --- |
| ANK repeat | 4 | 3.60E-02 |
| Ankyrin repeat | 4 | 4.70E-02 |

Cluster 6 **enrichment score: 1.33**
|  | Count | *p* value |
| --- | --- | --- |
| Carbohydrate metabolic process | 5 | 1.40E-02 |

Gene prediction was conducted with Augustus [52] on the scaffolds of the *D. guttifera* inbred line (A5). The option used in the analysis with Augustus was “-- species=fly”.

### Collecting samples for transcriptome analysis

Pupae at stage P12 (i) or P12 (ii) were used for two successive transcriptome analyses. These stages are just after the stage when *yellow* expression and pigmentation process have started [53]. The transcriptomes were compared by performing the combination of utilization of fluorescence-marked tissue, repetitive microsurgical samplings, and a sensitive RNA sequencing technology. For the first experiment, individuals from the transgenic line No. 1 which carries *eGFP* connected with an enhancer of *yellow* were used. An EGFP positive area around a campaniform sensillum on 3rd longitudinal vein (Area 1, Fig. S1a), an EGFP positive area at the tip of 3rd longitudinal vein (Area 2, Fig. S1a), and an EGFP negative area on 3rd longitudinal vein in wings of flies (Area 3, Fig. S1b) were separated with a surgical knife under a stereo microscope SZX-16 (Olympus). The width of these tissues was about 50 μm For the second experiment, an EGFP-positive area and an EGFP-negative area with a width of about 75 μm at the same place in individuals from transgenic lines No. 1 and No. 2 were dissected (Fig. S1c, d). From dissected tissues, RNA was collected with an RNeasy Micro Kit (Qiagen) and stored at −80°C. For RNA extraction, 20 dissected tissues were used for one replicate. Five biological replicates for each area were prepared (total: 20 × 5 = 100 tissues). The quality of extracted RNA was examined with an Agilent 2100 bioanalyzer (Agilent Technology).

### RNA sequencing

The library for RNA sequencing was constructed according to the protocol of Quartz-Seq, a highly sensitive method of RNA sequencing [54]. This protocol includes two PCR steps. Twenty-one cycles were performed for the first PCR, and eight cycles were performed for the second PCR. RNA sequencing was performed with NextSeq 550 (Illumina).

### Transcriptome analysis and enrichment analysis

The sequenced transcriptome was mapped to the genome of *D. guttifera* with HISAT2 [55]. Transcriptome assembly was conducted with StringTie [56]. Differentially expressed genes were identified with edgeR [57]. An FDR (false discovery rate) of 0.05 was chosen as the threshold to identify DEGs.

DEGs were blasted against the protein database of *D. melanogaster*, obtained from Ensembl [58,59]. BLAST analysis was performed with Blastx using an E-value < 1e-3. The top hit outcome for each gene was taken as the result of gene annotation. Based on the obtained gene annotation, enrichment analysis was conducted with DAVID [60]. Genes that could not be annotated was reanalyzed with Blast2GO (database: nr, E-value < 1e-3) [61].

## Authors’ contributions

Y.F., S.Kondo, S.S. and S.Koshikawa conceived of and designed the study; Y.F., S.Kondo, and A.T. collected data and conducted the analysis; Y.F., A.T. and S.Koshikawa drafted the manuscript, with input from S.Kondo and S.S.; Y.F. and S.Koshikawa edited and wrote the final version; all authors gave final approval for publication and agree to be held accountable for the work performed therein.

## Data accessibility

Sequence reads were submitted to the DDBJ/EMBL/GenBank. BioProject: PRJDB9109, BioSample: SAMD00198200-SAMD00198299. PE400 (DRX055245); MP3000 (DRX055246); MP5000 (DRX055247).

## Acknowledgements

We thank Sean B. Carroll and Thomas Werner for providing fly stocks; Tsuyoshi Katahata and Machiko Teramoto for fly stock maintenance; Kiyokazu Agata, Noriko Funayama, Keiji Matsumoto, Naoyuki Fuse, Takeshi Inoue, Norito Shibata, Katsushi Yamaguchi, and Takahiro Bino for scientific advice; and Elizabeth Nakajima for English editing. This work was supported by Functional Genomics Facility, NIBB Core Research Facilities.

## Supplementary figure and table legends

**Fig. S1.**
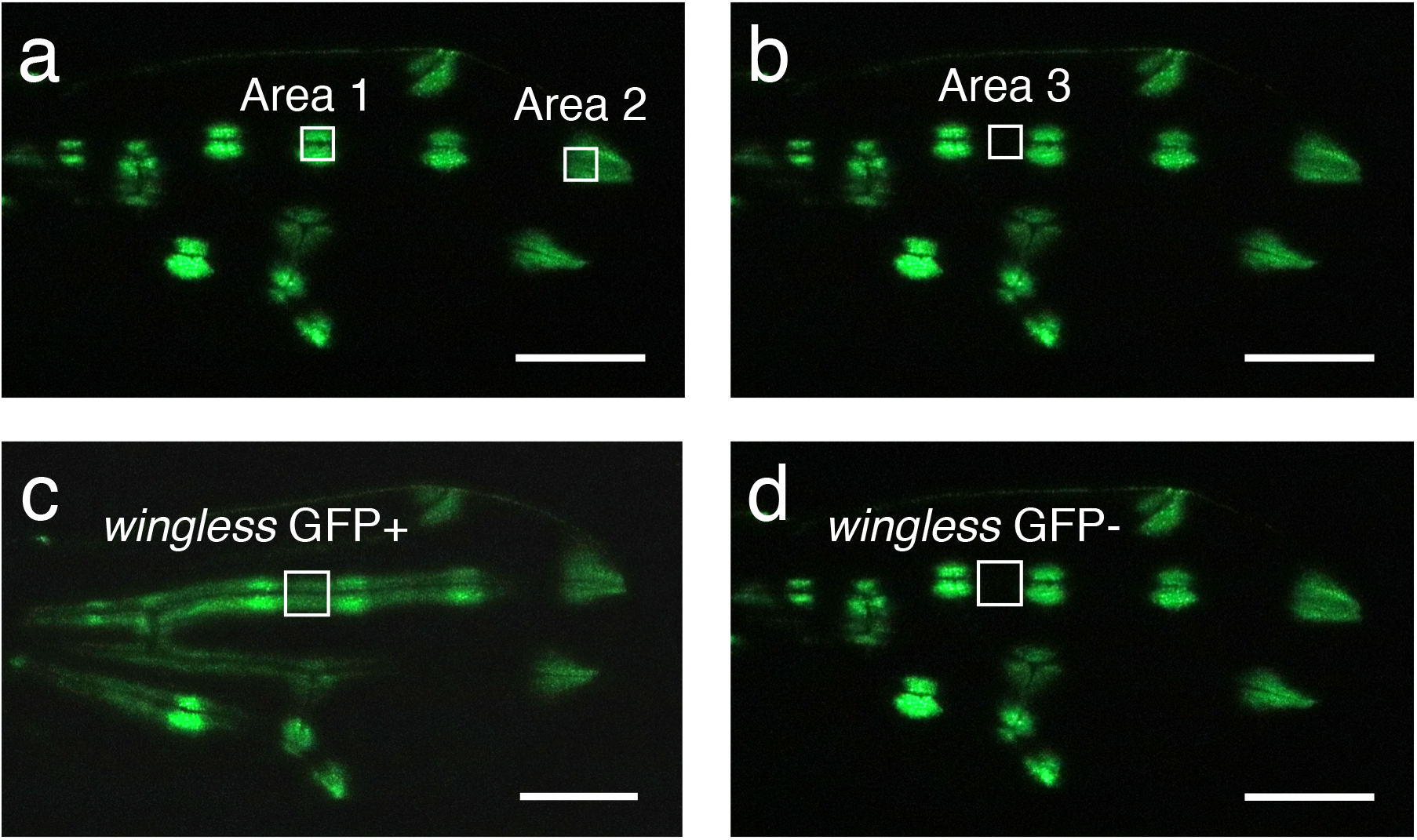
Tissues dissected for transcriptome analyses are shown. Dissected areas are indicated with white boxes. **a**: Area 1 (an EGFP positive area around a campaniform sensillum on 3rd longitudinal vein) and Area 2 (an EGFP positive area at the tip of 3rd longitudinal vein). **b**: Area 3 (an EGFP negative area on 3rd longitudinal vein). **c**: An EGFP positive area where *wingless* is ectopically expressed. **d**: An EGFP negative area where *wingless* is not ectopically expressed. Scale bars indicate 250 μm

**Table S1.** Genes differentially expressed in a pigmentation area around a campaniform sensillum. FC, CPM, and FDR indicate fold change, counts per million, and false discovery rate, respectively.

|  | logFC | logCPM | FDR |
| --- | --- | --- | --- |
| *y* | 4.97989524 | 13.1819431 | 2.38E-36 |
| *t* | 2.76380354 | 12.0058893 | 3.29E-32 |
| *laccase2* | 2.49456691 | 11.8424645 | 2.15E-33 |
| *yellow-e* | -1.6407385 | 10.3188765 | 1.81E-15 |
| *yellowe-h* | -0.5731037 | 10.5879713 | 0.02343591 |
| *svr* | -0.7948767 | 9.41379963 | 0.00438384 |
| *ZIPIC* | 4.53189996 | 0.48568358 | 0.0208437 |
| *zinc finger*<br>*protein 28-like* | 6.75852343 | 1.85690936 | 0.0005392 |
| *E(spl)m3-HLH* | 1.23731928 | 8.47601255 | 0.0003062 |
| *E(spl)m5-HLH* | 6.77199195 | 3.84590398 | 6.05E-10 |
| *E(spl)m7-HLH* | 3.36253299 | 4.92651899 | 0.0000107 |
| *Mad* | -1.5333172 | 5.94437046 | 0.02335105 |
| *DI* | 1.70066765 | 8.69275453 | 3.05E-07 |
| *Wnt4* | 7.88039218 | 2.78881957 | 0.00610713 |
| *sax* | 1.84041069 | 5.74292161 | 0.04887547 |
| *fz2* | -1.7268879 | 8.84471432 | 1.56E-08 |
| *wg* | 4.90156885 | 0.66703408 | 0.00181758 |

**Table S2.**
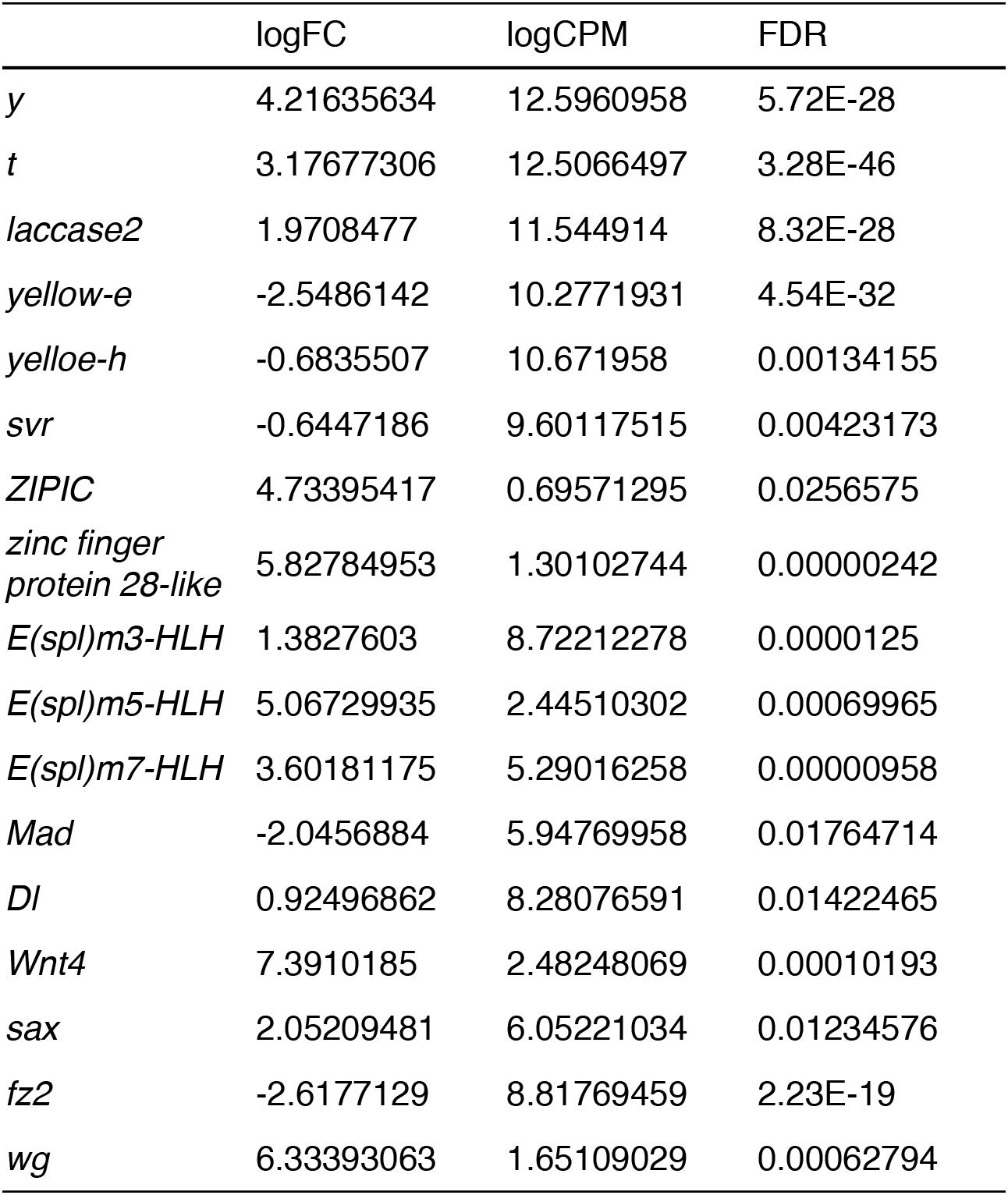
Genes differentially expressed in a pigmentation area at the tip of a vein. FC, CPM, and FDR indicate fold change, counts per million, and false discovery rate, respectively.

|  | logFC | logCPM | FDR |
| --- | --- | --- | --- |
| *y* | 4.21635634 | 12.5960958 | 5.72E-28 |
| *t* | 3.17677306 | 12.5066497 | 3.28E-46 |
| *laccase2* | 1.9708477 | 11.544914 | 8.32E-28 |
| *yellow-e* | -2.5486142 | 10.2771931 | 4.54E-32 |
| *yellowe-h* | -0.6835507 | 10.671958 | 0.00134155 |
| *svr* | -0.6447186 | 9.60117515 | 0.00423173 |
| *ZIPIC* | 4.73395417 | 0.69571295 | 0.0256575 |
| *zinc finger*<br>*protein 28-like* | 5.82784953 | 1.30102744 | 0.00000242 |
| *E(spl)m3-HLH* | 1.3827603 | 8.72212278 | 0.0000125 |
| *E(spl)m5-HLH* | 5.06729935 | 2.44510302 | 0.00069965 |
| *E(spl)m7-HLH* | 3.60181175 | 5.29016258 | 0.00000958 |
| *Mad* | -2.0456884 | 5.94769958 | 0.01764714 |
| *DI* | 0.92496862 | 8.28076591 | 0.01422465 |
| *Wnt4* | 7.3910185 | 2.48248069 | 0.00010193 |
| *sax* | 2.05209481 | 6.05221034 | 0.01234576 |
| *fz2* | -2.6177129 | 8.81769459 | 2.23E-19 |
| *wg* | 6.33393063 | 1.65109029 | 0.00062794 |

**Table S3.** Genes differentially expressed in a area where *wingless* is ectopically expressed. FC, CPM, and FDR indicate fold change, counts per million, and false discovery rate, respectively.

|  | logFC | logCPM | FDR |
| --- | --- | --- | --- |
| *y* | 4.54116336 | 12.8494004 | 3.6E-148 |
| *t* | 2.40367879 | 12.0035577 | 1.25E-74 |
| *laccase2* | 2.04478128 | 11.4946868 | 3.67E-39 |
| *yellow-e* | -1.2740677 | 11.392683 | 1.96E-25 |
| *yellowe-h* | -0.4516609 | 9.7843299 | 0.00750851 |
| *svr* | -0.6215228 | 9.01017126 | 0.00382779 |
| *ZIPIC* | 5.0147075 | -0.3005216 | 0.0284088 |
| *zinc finger*<br>*protein 28-like* | 5.61711109 | 2.5793453 | 0.00012374 |
| *E(spl)m3-HLH* | 0.53145394 | 8.24183597 | 0.00380777 |
| *E(spl)m5-HLH* | 5.25665975 | 1.58014564 | 0.00043059 |
| *E(spl)m7-HLH* | 1.67127368 | 5.65074921 | 1.23E-15 |
| *Mad* | -0.8277961 | 4.89916451 | 0.01393637 |
| *DI* | 1.02257261 | 7.82859549 | 8.12E-11 |
| *Wnt4* | 8.32686641 | 2.16749637 | 4.35E-09 |
| *sax* | 0.91143441 | 5.58205955 | 0.00742031 |
| *fz2* | -0.4058316 | 8.27179547 | 0.02026252 |
| *wg* | 0 | -1.2863604 | 1 |

**Table S4.**
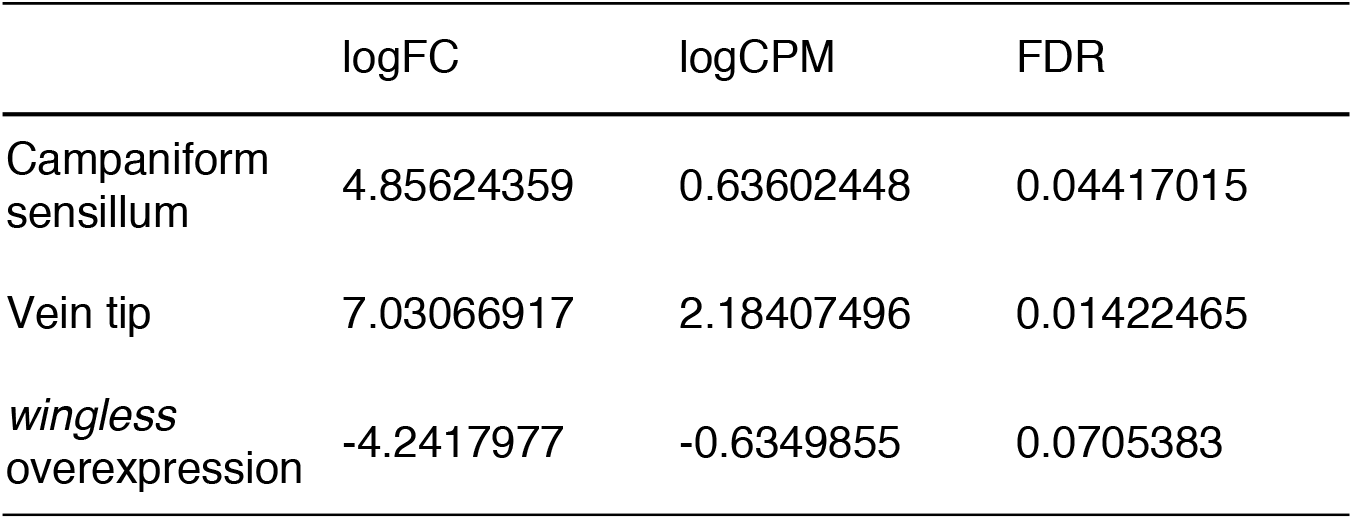
Data of expression of *hedgehog* gene in a pigmentation area around a campaniform sensillum, at the tip of a vein, and in a area where *wingless* is ectopically expressed.

